# CancerFoundation: A single-cell RNA sequencing foundation model to decipher drug resistance in cancer

**DOI:** 10.1101/2024.11.01.621087

**Authors:** Alexander Theus, Florian Barkmann, David Wissel, Valentina Boeva

## Abstract

We present CancerFoundation, a novel single-cell RNA-seq foundation model (scFM) trained exclusively on malignant cells. Despite being trained on only one million total cells, a fraction of the data used by existing models, CancerFoundation outperforms other scFMs in key tasks such as zero-shot batch integration and drug response prediction. During training, we employ tissue and technology-aware oversampling and domain-invariant training to enhance performance on underrepresented cancer types and sequencing technologies. We propose survival prediction as a new downstream task to evaluate the generalizability of single-cell foundation models to bulk RNA data and their applicability to patient stratification. CancerFoundation demonstrates superior batch integration performance and shows significant improvements in predicting drug responses for both unseen cell lines and drugs. These results highlight the potential of focused, smaller foundation models in advancing drug discovery and our understanding of cancer biology. Our code is available here^1^.

## 1 Introduction

Cancer is a leading cause of death globally, with millions of fatalities annually (Siegel et al., 2024). Despite extensive research and advancements in treatment, curing cancer remains challenging, largely due to the heterogeneity within tumor tissues. Tumors comprise various malignant and non-malignant cells, each exhibiting diverse genetic profiles and transcriptional states, influencing tumor progression, metastasis, and treatment resistance (Vasan et al., 2019; McGranahan et al., 2015). Understanding the biology of cancer transcriptional states and the cellular plasticity between the states is crucial, as these factors impact the tumor’s behavior and response to therapies. Indeed, previous research has linked cancer states to key cancer characteristics such as stemness, proliferation, and drug resistance across various cancer types (Neftel et al., 2019; Kim et al., 2020).

Single-cell RNA sequencing (scRNA-seq) has revolutionized the study of tumor heterogeneity, allowing for detailed mapping of gene expression in individual cells (Sikkema et al., 2023). Machine learning techniques have increasingly been applied to scRNA-seq data to uncover complex patterns and predict cellular behavior. Notably, the development of foundation models, originally conceived in the realm of natural language processing (NLP), has begun to influence this field. Foundation models are large-scale, pre-trained models that have demonstrated remarkable capabilities across various NLP tasks, such as translation, summarization, and question-answering (Brown et al., 2020a). Their success has prompted researchers to extend their application to other data modalities, including images (Dosovitskiy et al., 2021), proteins (Rao et al., 2021), and as of recent, scRNA-seq data (Cui et al., 2023; Hao et al., 2023; Rosen et al., 2023).

However, current single-cell foundation models (scFMs) have significant limitations when studying cancer biology. In particular, they are mainly trained on non-malignant cells. This limits their ability to accurately represent the complexities of tumors. For example, Copy-number alterations (CNAs), *i*.*e*., alterations in the copy number of specific genes or chromosomal regions, arise in malignant cells during cancer progression and treatment evasion and are closely associated with cancer transcriptional states (França et al., 2022). However, they are very rare in non-malignant cells.

To address these gaps, we propose *CancerFoundation*, a foundation model tailored specifically for scRNA-seq data from malignant cells, designed to capture the unique transcriptional states of cancer. Our aim is to create a model that not only provides a nuanced representation of tumor heterogeneity but also translates these representations into practical applications for drug discovery.

**The main contributions** of our work are:

i. We introduce CancerFoundation, a cancer-specific scRNA-seq foundation model trained only on malignant cells. CancerFoundation outperforms existing scFMs in integration and drug response prediction tasks with up to 10 times fewer parameters and 50 times less training data.
ii. We demonstrate that tissue and technology-aware oversampling enhances performance on underrepresented tissues and technologies.
iii. We propose survival prediction as a novel downstream task to evaluate the generalizability of scFMs to bulk RNA data and their applicability to patient stratification.

## 2 Background

A promising approach for applying machine learning to scRNA-seq data is the development of foundation models, which leverage generative pre-training to achieve remarkable performance across various domains. These models are typically built on transformers, a deep learning architecture that allows them to process large-scale and diverse datasets efficiently. The most notable applications of foundation models can be seen in fields like computer vision and natural language generation (NLG), with models such as GPT-4 and DALL-E2 leading the way. These models demonstrate the potential of pre-training on extensive datasets, enabling them to be easily customized for a variety of downstream tasks.

In recent years, substantial efforts have been dedicated to extending foundation models beyond text and image data to encompass a wide range of data modalities, including audio (Yang et al., 2023), brain signals (Caro et al., 2024), and, of particular relevance to our work, scRNA-seq (Cui et al., 2023; Rosen et al., 2023; Hao et al., 2023). Among these, Universal Cell Embeddings (UCE) (Rosen et al., 2023), scGPT (Cui et al., 2023), and scFoundation (Hao et al., 2023) stand out as pioneering approaches. These models are designed to handle the complex, high-dimensional data generated by scRNA-seq, providing frameworks for integrating and interpreting single-cell datasets across different conditions and experimental settings.

**UCE**, for instance, aims to create a unified representation of cells from diverse tissues and species, facilitating cross-study comparisons and transfer learning. UCE abstracts cells as “bags of RNA” by transforming the RNA gene expression of a single cell into a sample of its corresponding genes, weighted by their expression levels. However, UCE only predicts whether a gene has been expressed or not, without providing quantitative details on the expression levels. Furthermore, genes are represented by their protein product using a large protein language model (Lin et al., 2023), enabling generalization across species. With 650 million parameters, UCE is at the time of this writing the largest foundation model for scRNA-seq.

In contrast, **scGPT** leverages the power of generative pre-trained transformers (Radford & Narasimhan, 2018; Radford et al., 2019; Brown et al., 2020b) to model gene expression profiles, specifically focusing on human cells. Unlike UCE, scGPT trains only on expressed genes and is designed to predict the (binned) expression values of these genes, providing more granular information about gene expression levels. Additionally, scGPT includes a separate model trained on a dataset of five million cancer cells, comprising both malignant and non-malignant cells.

**scFoundation** takes a different approach, employing an asymmetric transformer-like architecture with 100 million parameters trained on over 50 million cells, one fourth being tumor cells. Its pre-training process focuses on gene expression enhancement and harmonization across varying sequencing depths. scFoundation defines two total count indicators—”T” (target) and “S” (source)—which correspond to the total counts of the raw and input samples, respectively. During pre-training, the model predicts masked gene expression values, capturing gene-gene relationships within the cell and normalizing across different read depths. At inference, scFoundation can enhance sequencing depth by feeding the raw gene expression to the model and setting the “T” count higher than the original “S”, generating an enhanced gene expression profile. Notably, scFoundation demonstrates state-of-the-art performance in predicting anti-cancer drug responses.

To build upon these existing frameworks, we propose a foundation model specifically designed to address the challenges posed by cancer heterogeneity and drug response prediction. Unlike UCE, scGPT, and scFoundation, which generalize across various cell types and conditions, our method focuses exclusively on malignant cells, enabling a more targeted understanding of cancer-specific transcriptional programs. By concentrating solely on malignant cells, we aim to enhance the model’s ability to capture key biological patterns linked to tumor progression and drug resistance. Additionally, we incorporate data balancing techniques to mitigate the inherent biases present in malignant scRNA-seq data, ensuring that underrepresented populations are accurately modeled.

## 3 Methodology

### 3.1 Dataset

We curated an extensive dataset of one million high-quality malignant cells from the Curated Cancer Cell Atlas (Gavish et al., 2023). The dataset spans roughly 1,500 individual tumors from over 112 studies and 46 cancer types. We only included samples that contained at least 50 malignant cells and removed studies with fewer than four samples. Furthermore, InferCNV (Tickle et al., 2019) was used to infer the CNA status of each gene in each cell. The inferred CNA profiles were then used to better distinguish between malignant and non-malignant cells.

The dataset is highly imbalanced along various dimensions. In Figure 1, we visualize the imbalance for anatomic sites and scRNA-seq technologies. In particular, substantial discrepancies exist along anatomic sites (see Figure 1b). To highlight this point, breast cancer cells make up 20.4% of the entire dataset, while liver and biliary cancer cells only account for 0.43%. This imbalance is in direct conflict with our overarching objective to be *pan-cancer*, as training with the dataset as is would heavily bias us towards certain cancer types like breast cancer. Another significant imbalance commonly observed in scRNA-seq datasets is related to sequencing technologies (see Figure 1a), with 10x Genomics (Zheng et al., 2017) accounting for 87.9% of all cells examined. This imbalance is critical because the choice of technology directly affects the transcriptomic profiles obtained. In addition, different technologies are better suited to specific tissue types.

**Figure 1:**
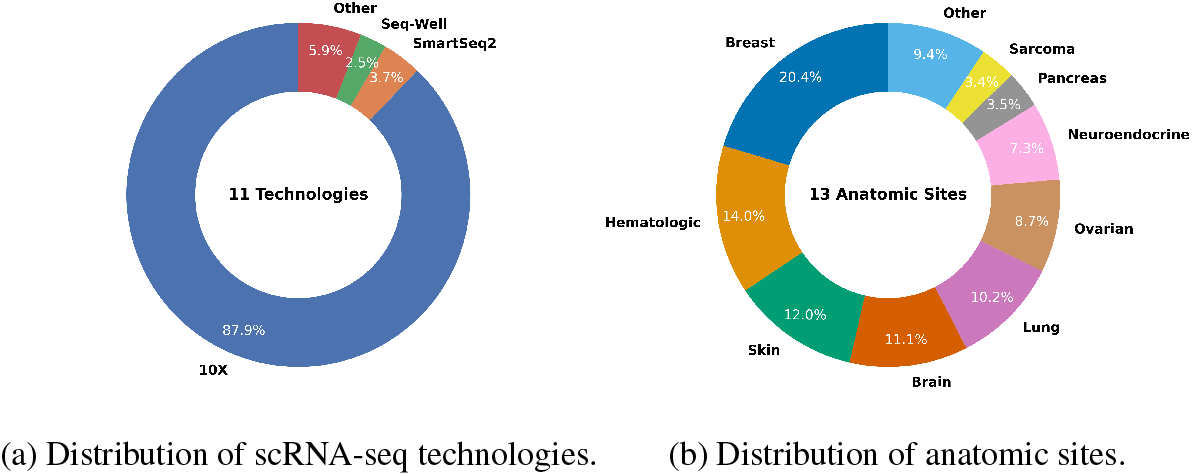
Distribution of scRNA-seq technologies (a) and anatomic sites (b) in the training dataset. The percentages indicate the proportion of cells within the training dataset.

For example, 10x Genomics excels at processing tissues that are easy to dissociate and produce large numbers of cells, such as blood and skin. In fact, 97.4% of the skin cancer cells in our dataset were collected using 10x Genomics. However, 10x Genomics struggles with dense or fragile tissues, such as the brain, which are harder to dissociate. These tissues often benefit from more sensitive methods like SmartSeq2 (Picelli et al., 2013), which, although lower in throughput and more expensive, provides full-length transcript coverage and tolerates partial dissociation. This makes SmartSeq2 better suited for capturing subtle changes in gene expression in tissues such as the brain. Indeed, 22.5% of the brain cancer cells in our dataset were processed with SmartSeq2.

Hence, the technology imbalance not only skews representation of different platforms but also limits performance in tissues that are harder to dissociate. Addressing this disparity is thus crucial to uphold our pan-cancer objective.

### 3.2 CancerFoundation

#### 3.2.1 Data sampling

##### Cell-based sampling

As discussed in Section 3.1, scRNA-seq dataset suffer from significant imbalances, both in anatomical sites and scRNA-seq technologies. Alsabbagh et al. (2023) recently investigated the impact of cell type imbalances on training scFMs. Predictably, they observed poor performance on underrepresented cell types. To address this, the authors proposed three sampling techniques and concluded that *random oversampling* performed best. This method involves increasing the representation of minority classes by sampling their data points multiple times within a training epoch.

In our case, however, a more nuanced approach is necessary due to the extreme nature of the imbalances—especially in terms of the technologies, as shown in Figure 1a—and the fact that we are dealing with two independent dimensions of imbalance (technology and anatomic site). In particular, we apply a stratified oversampling strategy. Oversampling all minority sites to match the majority would lead to extreme sampling ratios (e.g., liver cells by a factor of 46), risking overfitting. Instead, sites are oversampled to tiers of 20%, 40%, and so on, reducing imbalance while maintaining data integrity. For scRNA-seq technologies, which are often linked to specific tissue types (e.g., 10x Genomics in skin, SmartSeq2 in brain), we prioritize balancing minority technologies at each site, capping their oversampling factor at 10 to prevent overfitting. This approach reduces imbalances without distorting natural variations across tissues.

##### Gene-based sampling

For a cell *i*, let 𝔾^(*i*)^ be its set of genes with cardinality *N*^(*i*)^, and *M* the maximum sequence length, where *M, N*^(*i*)^ ∈ ℕ^+^ and *M* < *N*^(*i*)^. Models like scGPT randomly select *M* expressed genes (without replacement) from the set of expressed genes 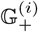:

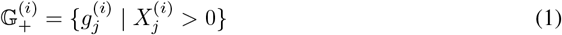

This approach has two key limitations: (1) High sparsity in scRNA-seq data limits the pool of expressed genes, potentially biasing the model toward a small set of highly expressed genes, and (2) it overlooks non-expressed genes, which contain important biological information about cell identity and tissue specificity.

To address this, we propose a hybrid method that selects a mix of expressed and non-expressed genes. A proportion of expressed genes is selected from 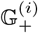, and the remainder from the set of non-expressed genes, 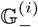:

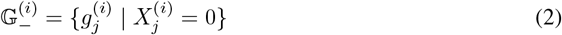

This ensures a balance between expressed and non-expressed genes, enhancing diversity in the model input while preserving important biological signals.

#### 3.2.2 Input embeddings

Integrating scRNA-seq data into CancerFoundation is challenging due to the presence of both categorical (genes) and continuous elements (expression values), unlike the text data used in standard GPT models.

##### Gene tokens

Similar to existing scFMs, we treat genes as fundamental units of information, assigning each a unique integer identifier, along with special tokens `<cls>` and `<pad>` for cell-level context and padding, respectively. The gene sequence for each cell has a fixed length *M*, represented as:

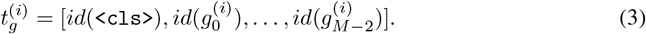

To reduce complexity, we focus on a subset of 29 thousand highly variable genes relevant to each tissue type, rather than using all 67 thousand genes, as in scGPT, which improves computational efficiency and model interpretability.

##### Expression values

Expression values vary significantly between sequencing methods, making standard preprocessing insufficient. To standardize across cells, we use scGPT’s binning technique, transforming continuous expression values into discrete categories. This reduces biases and harmonizes the data across batches, enhancing the model’s robustness.

##### Final input

To obtain the final input for CancerFoundation, we use the gene tokens 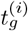 to index a conventional embedding layer^1^, denoted as emb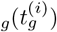. In order to preserve the ordinal nature of the binned expression values 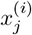, we pass them to a two-layer multi-layer perceptron (MLP). The embedding for the gene tokens, and the binned expression values is finally summed up to obtain the final input to our model, *h*^(*i*)^:

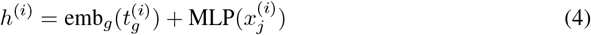

#### 3.2.3 Model

We utilize a relatively compact self-attention transformer model based on the architecture detailed by Vaswani et al. (2023), though it is implemented without positional encoding. Our model consists of six transformer layers, an embedding dimension of 256, and a hidden dimension of 512. This design yields a significantly smaller model compared to other leading foundation models for scRNA-seq data. Namely, our model has only 10.8 million parameters, making it over 60 times smaller than UCE (Rosen et al., 2023), 11 times smaller than scFoundation (Hao et al., 2023), and 5 times smaller than scGPT (Cui et al., 2023). This substantial reduction in size is achieved through our novel strategy that focuses on training with a much smaller dataset of malignant cells, as opposed to the larger datasets of 30-50 million cells, primarily composed of peripheral blood mononuclear cells, used by other models.

#### 3.2.4 Pretraining objective

Self-attention effectively captures complex dependencies in sequential data like natural language. There are two prominent strategies: (1) masked token prediction, as used in BERT (Devlin et al., 2019) and Roberta (Liu et al., 2019), where tokens are masked and predicted by the model; and (2) autoregressive generation in models like GPT (Radford & Narasimhan, 2018; Radford et al., 2019; Brown et al., 2020b), which predicts tokens sequentially. Masked token prediction models leverage bidirectional attention, whereas autoregressive models apply unidirectional attention.

In our work, we use a hybrid approach to adapt these methods for gene expression data, which lacks a strict sequential order. We employ masked token prediction but modify the attention mechanism so that non-masked tokens cannot attend to masked tokens, thereby combining generative behavior without enforcing a unidirectional sequence constraint.

For training, we randomly mask 25%, 50%, or 75% of the gene expression values, replacing them with a placeholder (e.g., −1). While this setup resembles scGPT, our model introduces a key modification in loss calculation using domain-invariant training. Given a set of masked genes *U*_*unk*_, the output for cell *i* and gene *n*, 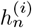, is concatenated with a technology-specific embedding and passed through a multi-layer perceptron (MLP) to predict expression values using mean squared error:

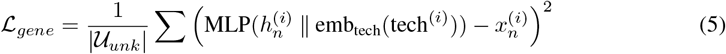

Additionally, to capture global cell information, a `<cls>` token is trained with a similar objective:

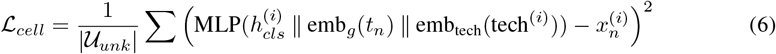

By integrating technology-specific embeddings, our model aligns cross-domain representations, enhancing performance on minority technologies.

## 4 Results

### 4.1 Zero-shot batch integration

Batch integration in cancer scRNA-seq data is challenging, as malignant cells often cluster by patient rather than by transcriptional state. To address this, we evaluated CancerFoundation’s ability to perform zero-shot integration — integrating new datasets that the model has neither seen during pretraining nor been finetuned on. This capability is essential for downstream tasks like drug response prediction, where the model must generalize to unseen datasets.

We compared CancerFoundation’s integration performance against two leading models: scVI (Lopez et al., 2018) and scGPT (Cui et al., 2023). scVI, a variational autoencoder, is known for robust batch correction but is not a foundation model and requires training on the test dataset. In contrast, scGPT and CancerFoundation are transformer-based foundation models. The cancer-specific version of scGPT, denoted as “scGPT (cancer)”, was pretrained on malignant and non-malignant cells, while CancerFoundation is pretrained exclusively on malignant ones. We assessed these models using the framework from Luecken et al. (2022).

This comparison highlights CancerFoundation’s unique focus on malignant cell states, potentially enhancing its ability to preserve biological accuracy across batches. While scGPT (cancer) benefits from broader cell type exposure, CancerFoundation’s specialization may offer advantages in recognizing subtle transcriptional differences in cancer cells.

We validated CancerFoundation’s performance using the glioblastoma dataset from Neftel et al. (2019), which includes four distinct malignant cell states: neural-progenitor-like (NPC-like), oligodendrocyte-progenitor-like (OPC-like), astrocyte-like (AC-like), and mesenchymal-like (MES-like). Glioblastoma is a highly heterogeneous brain tumor, making it an ideal test case for assessing batch integration in complex transcriptional landscapes. Refer to Appendix C for further results, including an ablation study highlighting the effect of our data balancing strategies.

CancerFoundation significantly outperformed both scVI and scGPT, as well as its cancer-specific variant, in preserving the integrity of biological signals across batches, as shown in Table 1. Following CancerFoundation, the scGPT (cancer) variant demonstrated slightly better biological conservation metrics compared to standard scGPT, particularly in terms of the cell-type local inverse Simpson index (cLISI) (Korsunsky et al., 2019) metric. Batch correction metrics were similar across the three foundation models, with scGPT showing a slight advantage, primarily due to the k-nearest-neighbor batch-effect (kBET) metric (Büttner et al., 2019). The only non-foundation model, scVI, was a clear outlier, performing worst in preserving transcriptional states, while achieving superior batch correction results, particularly in the kBET score.

**Table 1:** Batch integration metrics evaluating the performance of different embeddings in integrating the batches for a glioblastoma dataset (Neftel et al., 2019) using the framework introduced by Luecken et al. (2022).

| Method | Biological conservation |  |  |  |  | Batch correction |  |  |  | Aggregate score |  |  |
| --- | --- | --- | --- | --- | --- | --- | --- | --- | --- | --- | --- | --- |
|  | Isolated labels | Leiden NMI | Leiden ARI | Silhouette label | cLISI | Silhouette batch | iLISI | KBET | GC | Batch correction | Biological conservation | Total |
| CancerFoundation (ours) | **0.59** | **0.51** | **0.48** | **0.55** | **0.92** | 0.87 | 0.07 | 0.26 | **0.93** | 0.53 | **0.61** | **0.58** |
| scGPT (cancer) | 0.57 | 0.40 | 0.31 | 0.53 | 0.89 | **0.88** | 0.07 | 0.28 | **0.93** | 0.54 | 0.54 | 0.54 |
| scGPT | 0.55 | 0.39 | 0.31 | 0.52 | 0.83 | 0.87 | 0.10 | 0.35 | **0.93** | 0.56 | 0.52 | 0.54 |
| scVI | 0.54 | 0.29 | 0.20 | 0.51 | 0.70 | 0.84 | **0.20** | **0.61** | 0.92 | **0.64** | 0.45 | 0.47 |

Figure 2 presents UMAP (McInnes et al., 2020) representations of the embeddings, which corroborate the findings derived from the metrics. scVI appears to overcorrect for batch effects, resulting in blurred distinctions between transcriptional states. The scGPT (cancer) model shows an improvement over scVI, but struggles to clearly separate MES-like cells from AC-like ones. In contrast, our model, CancerFoundation, distinguishes all four states effectively.

**Figure 2:**
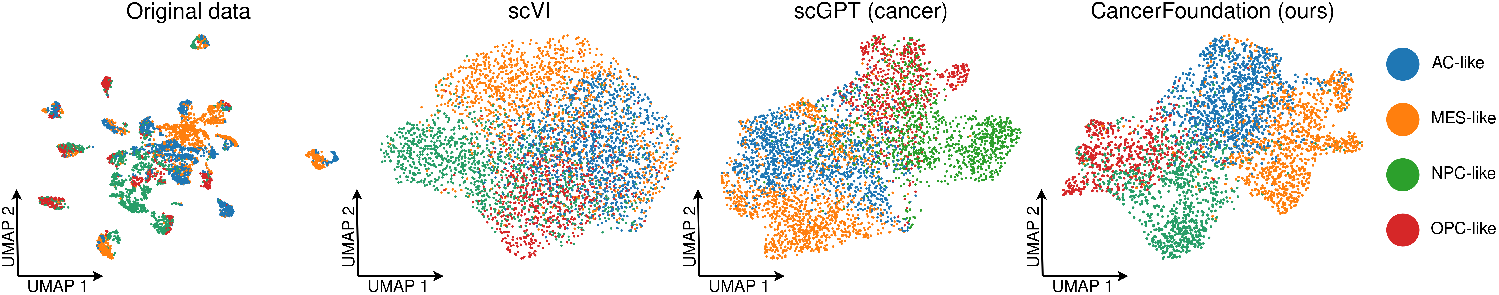
UMAP plots of cell embeddings colored according to the ground-truth transcriptional states for a glioblastoma dataset (Neftel et al., 2019): AC-like (astrocyte-like, blue), MES-like (mesenchymal-like, yellow), NPC-like (neural-progenitor-like, green), and OPC-like (oligodendrocyte-progenitor-like, red).

### 4.2 Drug response prediction

Cancer Drug Responses (CDRs) assess how tumor cells react to drug treatments. Accurately predicting CDRs through computational methods is crucial for developing anti-cancer therapies and gaining insights into cancer biology (Unger et al., 2015). Previousy, a foundation model for scRNA-seq data, scFoundation (Hao et al., 2023), integrated its model with a CDR prediction model called DeepCDR (Liu et al., 2020) to forecast the half-maximal inhibitory concentrations (IC_50_) of drugs across various cell line datasets. This experiment aimed to validate whether scFoundation could generate meaningful embeddings for bulk gene expression data, despite being trained on single-cell data.

The original DeepCDR model utilized drug structural information and multi-omics data to predict IC_50_ values. On the other hand, scFoundation focused on gene expression data and substituted the transcriptome multi-layer perceptron (MLP) subnetwork in DeepCDR with their foundation model. This allowed them to extract transcriptome features from scFoundation and feed them into the prediction module. They combined data from the Cancer Cell Line Encyclopedia (CCLE) (Barretina et al., 2012) and the Genomics of Cancer Drug Sensitivity (GDSC) (Iorio et al., 2016) databases to source cell line gene expression data, drugs, and IC_50_ values.

Despite being trained exclusively on single-cell rather than bulk gene expression data, scFoundation achieves significant improvements in accuracy over DeepCDR for both unseen cell lines and anticancer drugs, demonstrating an emergent ability.

In this section, we aim to assess whether our model, CancerFoundation, exhibits a similar ability and if it can surpass scFoundation’s performance. As discussed in Section 2, scFoundation was trained on a large dataset of 50 million cells, with roughly a quarter being tumor cells. In contrast, our model was trained on a much smaller dataset of only one million malignant cells. These experiments will help us explore whether focusing on a smaller, malignant-specific dataset offers advantages over training on a larger, more diverse scRNA-seq dataset.

#### 4.2.1 Hold-out cell lines

First, we predicted the treatment response IC_50_ values for the hold-out cell lines. Figure 3 shows that CancerFoundation outperforms DeepCDR across all cancer types, and almost all drugs by a large margin. Compared to scFoundation’s predictions, it exhibits comparable or superior accuracy for all cancer types except for stomach adenocarcinoma, a cancer type our foundation model was not trained on. In a similar manner, CancerFoundation significantly improves the predictions on the basis of drugs compared to scFoundation, especially in cases where scFoundation underperforms. For instance, the Pearson’s *R* for Lapatinib, an orally active drug used to treat breast cancer and other solid tumors (Burris, 2004), increased drastically from **0.08** to **0.95**.

**Figure 3:**
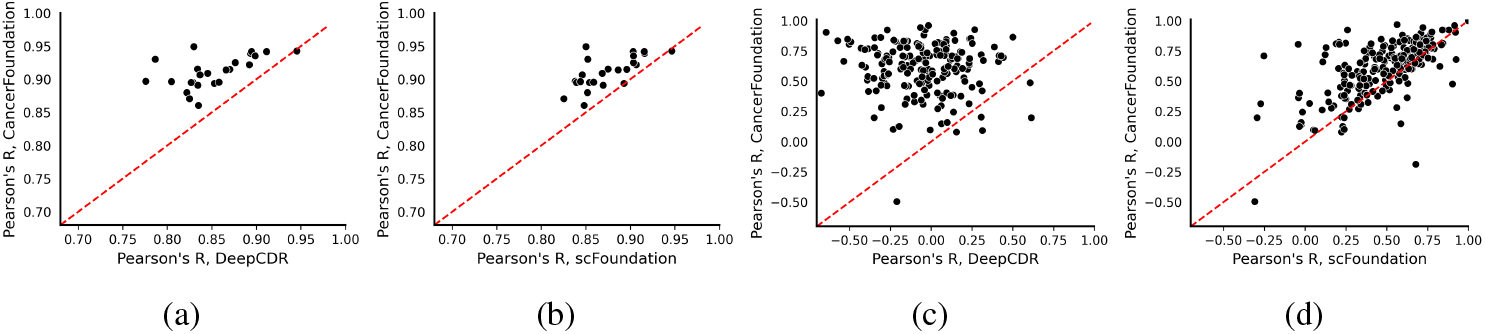
Comparison of Pearson’s *R* of predicted IC_50_ and measured IC_50_ for hold-out cell lines. Pearson’s *R* is computed across all drugs and cancer types in the test set. Each dot represents a cancer type (a, b) or drug (c, d). The x-axis shows Pearson’s *R* for DeepCDR (a, c) and scFoundation (b, d), respectively. The y-axis shows Pearson’s *R* from our model.

#### 4.2.2 Hold-out drugs

Lastly, to assess the generalization capabilities of our model, we evaluated its performance on hold-out drugs, which were excluded from the training set. It is important to note that the results for scFoundation deviate from those listed in their paper. In particular, they used early stopping based on validation performance using unseen drugs, which inadvertently introduced *data leakage*. By doing so, the model was indirectly exposed to information about the hold-out drugs during training, leading to inflated performance metrics. In contrast, we ensured that the unseen drugs were entirely excluded from the training and validation process, including early stopping, thus providing a more rigorous evaluation of the model’s generalization ability.

Figure 4 illustrates that CancerFoundation exhibites superior performance on hold-out drugs, achieving a higher Pearson’s *R* and notably lower standard deviation compared to both scFoundation and DeepCDR. While both scFoundation and CancerFoundation made reasonable predictions based on their Pearson’s *R* values, DeepCDR, which depends on raw bulk gene expressions, had difficulties generalizing effectively.

**Figure 4:**
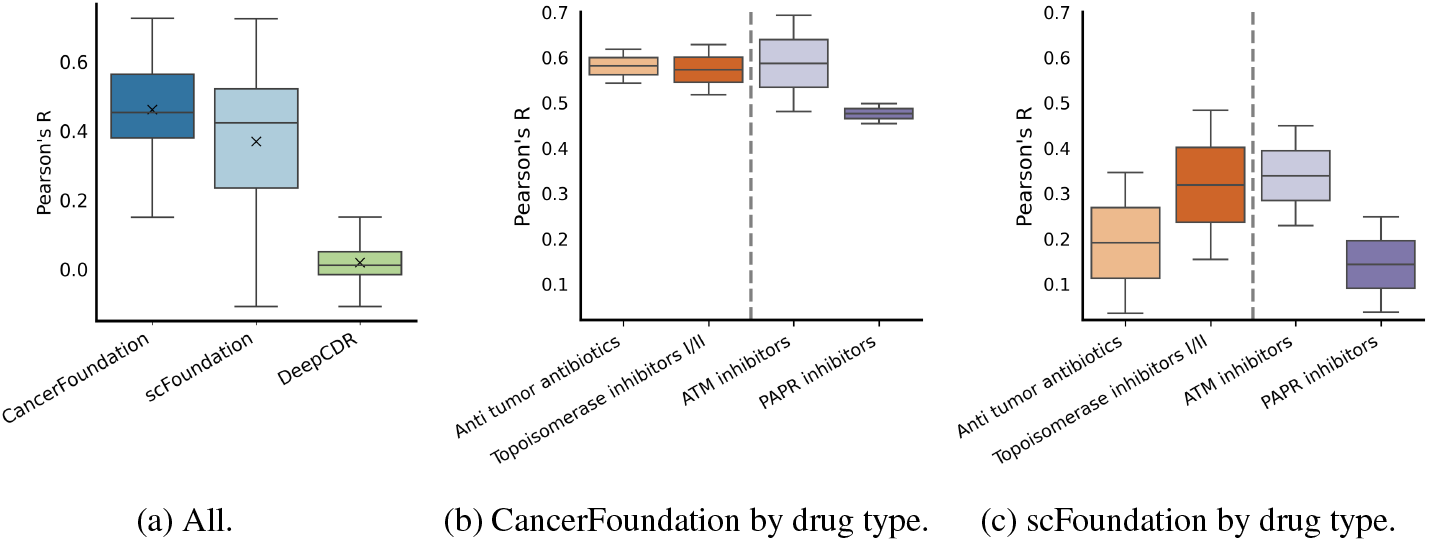
Comparison of Pearson’s *R* of predicted IC_50_ and measured IC_50_ for hold-out drugs across models and drug types. (a) Overall Pearson’s *R* comparison for CancerFoundation, scFoundation, and DeepCDR; the solid line represents the median and the cross represents the mean. (b) Pearson’s *R* comparison grouped by drug type for CancerFoundation: chemotherapy, anti-tumor antibiotics, and topoisomerase inhibitors I/II (left, orange); targeted therapy, PAPR inhibitors, and ATM inhibitors (right, purple). (c) Same drug type grouping for scFoundation.

Moreover, we categorized the drugs into various therapeutic classes to investigate if the accuracy of IC_50_ predictions was linked to their underlying mechanisms. The results of the scFoundation paper indicated that drugs used in chemotherapy, including anti-tumor antibiotics and topoisomerase inhibitors, exhibited a higher Pearson correlation coefficient (PCC) compared to those used in targeted therapy, such as ATM and PARP inhibitors. This discrepancy was explained by the fact that specific gene mutations significantly influence targeted therapies (Unger et al., 2015), but mutation details are challenging to extract from gene expression data. In contrast, chemotherapy drugs are often associated with gene expression patterns, making their IC_50_ predictions more straightforward. Given the concerns raised about data leakage in the previous evaluation, the validity of those results might be compromised. Indeed, when we correct for data leakage, the results indicate no significant difference between drugs used in chemotherapy and targeted therapy for scFoundation as shown in Figure 4c. As for CancerFoundation, predictions for drugs utilized in targeted therapy were overall worse than those used in chemotherapy, mainly driven by the low accuracy for PAPR (poly(ADP-ribose) polymerase) inhibitors compared to the other drug types (see Figure 4b).

### 4.3 Survival prediction

As a final and novel downstream task for scFMs, we propose to consider the patient stratification ability of embeddings generated on bulk RNA-seq data of large-scale cancer cohorts such as the PANCANATLAS of The Cancer Genome Atlas (TCGA) (Weinstein et al., 2013). In particular, we used the 21 cancer types selected by Wissel et al. (2022). For each cancer type, we selected the 1,199 most highly expressed genes to compute embeddings and to use as a baseline. We trained Ridge-regularized Cox Proportional Hazards (Cox PH) models on the embeddings generated by CancerFoundation (ours), scGPT (cancer), scGPT, and a baseline consisting of the same genes used for the embedding generation of all scFMs (Baseline (GEX)) (Cox, 1972; Breslow, 1975; Hoerl & Kennard, 1970; Benner et al., 2010). We measure the (discriminative) performance of all Cox PH models using Harrell’s concordance index (Harrell Jr et al., 1996).

Table 2 and Table 6 present the survival prediction performance of the Ridge-regularized Cox PH trained on different embeddings and gene expression as a baseline across different cancer types. The results, measured by Harrell’s concordance index, show that most scFMs do not significantly outperform the baseline trained on gene expression. However, for cancer types with substantial training data CancerFoundation often outperforms or matches the performance of both scGPT variants. This is particularly evident in GBM and LUSC, as shown in Table 2. Conversely, Table 6 demonstrates that CancerFoundation significantly underperforms the baseline for cancer types with little or no training data. This contrast highlights the importance of adequate training data for effectively applying CancerFoundation and suggests potential limitations in its generalizability to less-represented cancer types.

**Table 2:** Survival prediction performance for different embeddings and gene expression as baseline. Values show mean Harrell’s concordance index followed by standard deviation computed across 25 folds. Percentages in parentheses indicate the proportion of samples for each cancer type in the CancerFoundation training data. This table only shows the eight cancer types with the highest percentage of training data. See Table 6 for the remaining datasets.

| Method | BRCA (16%) | LUAD (8%) | GBM (7%) | PAAD (6%) | OV (6%) | SKCM (5%) | LAML (4%) | LUSC (4%) |
| --- | --- | --- | --- | --- | --- | --- | --- | --- |
| CancerFoundation (ours) | 0.56±0.05 | 0.59±0.06 | 0.59±0.07 | 0.51±0.08 | 0.51±0.05 | 0.52±0.13 | 0.60±0.07 | 0.56±0.06 |
| scGPT (cancer) | 0.54±0.06 | 0.61±0.06 | 0.53±0.04 | 0.49±0.09 | 0.52±0.06 | 0.63±0.14 | 0.60±0.07 | 0.52±0.06 |
| scGPT | 0.54±0.05 | 0.60±0.06 | 0.52±0.05 | 0.50±0.09 | 0.51±0.04 | 0.47±0.11 | 0.55±0.07 | 0.53±0.07 |
| Baseline (GEX) | 0.60±0.06 | 0.61±0.05 | 0.56±0.08 | 0.52±0.08 | 0.57±0.08 | 0.50±0.12 | 0.65±0.08 | 0.48±0.04 |

## 5 Conclusion

In conclusion, CancerFoundation demonstrates that a domain-specific foundation model trained exclusively on malignant cells can achieve state-of-the-art performance in key tasks such as batch integration and drug response prediction, while being significantly smaller than existing scFMs. With only 10.8 million parameters, up to 60 times fewer than comparable models, CancerFoundation showcases that focused, smaller scFMs can outperform larger models. Although scFMs generally do not surpass linear baselines for survival prediction across most datasets, CancerFoundation shows improvements over existing models for several datasets with sufficient single-cell training data.

## Code and Data Availability

An implementation of CancerFoundation and its weights are available at https://github.com/BoevaLab/CancerFoundation along with instructions to reproduce results.

The training data can be made available upon request.

## Acknowledgments

We thank Shuyang Fan for helpful discussions about training single-cell foundation models.

FB is supported by the Swiss National Science Foundation (SNSF) (grant number 205321 207931).

## Appendix

### A Implementation details

In this section, we outline the implementation choices covering the software frameworks, hardware configurations, and optimization strategies employed.

#### Framework

The entire model was implemented and trained using PyTorch (Paszke et al., 2019), which provided the flexibility and modularity necessary for deep learning research and experimentation. PyTorch’s dynamic computation graph and seamless GPU acceleration made it an ideal choice for both development and large-scale distributed training.

#### Sequence length

The model was trained with a maximum sequence length of 1200 tokens. This allowed the model to capture more context while avoiding the exponential increase in computational complexity and memory usage that typically arises from longer sequences. scGPT also uses a sequence length of 1200 tokens, thus presenting a conservative option.

#### Flash attention

In transformer-based models, the self-attention mechanism typically has quadratic complexity with respect to the sequence length (Vaswani et al., 2023). Given that the model operates with a maximum sequence length of 1200, this presents a significant challenge, as longer sequences can quickly lead to excessive memory consumption and slow training times.

To mitigate this, we employed Flash Attention (Dao et al., 2022), a more memory-efficient variant of the standard attention mechanism. Flash Attention optimizes both memory usage and computation during the attention step by reducing the intermediate memory allocations, allowing the model to handle longer sequences more effectively. This optimization was crucial for maintaining feasible memory consumption while training on large batches, without compromising on the model’s ability to capture long-range dependencies.

#### Mixed precision

To further optimize the training process, mixed precision training was employed. Computations were performed using both 16-bit (half precision) and 32-bit (single precision) floating point numbers. This not only improved the speed of training but also reduced memory consumption, enabling larger batch sizes to be used without running out of GPU memory.

#### Distributed training

We used synchronous data parallelism across eight GPUs, which allowed for a larger effective batch size and significantly sped up the training process. Each GPU processed a different subset of the data, with gradients synchronized at the end of each iteration to ensure consistent model updates.

For handling data during training, we used LaminDB (Lamin Labs, Inc., 2024), a distributed data management system that facilitated efficient data access across the eight GPUs used in training. LaminDB provided a smooth interface for managing data pipelines and eliminated potential I/O bottlenecks when scaling across multiple GPUs.

### B Training

This appendix details the training procedures and strategies employed in developing the model. Furthermore, we provide learning curves to illustrate the model’s performance during training.

#### Optimizer and learning rate scheduling

We use the Adam optimizer (Kingma & Ba, 2017) with a learning rate scheduling approach that linearly increases the learning rate from 0.0 to 10^−4^ over a warmup period of 10^3^ training steps, and subsequently decreases to 0.0 on a cosine schedule for the remaining steps^1^.

#### Gradient clipping

To ensure stable training of our foundation model, we applied gradient clipping with a uniform norm of 1.0 across all parameters. This technique is commonly used in deep learning models, particularly transformers, to prevent the exploding gradient problem, where gradients can grow excessively large and destabilize training. This is especially important when training with masked self-supervised learning, as the model may encounter complex input patterns that lead to volatile gradients (Devlin et al., 2019; Raffel et al., 2023).

#### Batch size

We selected a batch size of 64. The choice of batch size is a critical factor in this type of model, as it directly impacts both the computational efficiency and the quality of the learned representations. In our case, a batch size of 64 provided a good balance between stability during training and effective learning, especially given the large scale of the model and the self-supervised nature of the task.

#### Learning curves

In Figure 5, we present the evolution of both the training and evaluation losses. It is important to interpret the training losses with caution, as predicting gene expression often also involve batch effects. As discussed in Section 4.1, our goal is not to perfectly reconstruct the original data, but to capture transcriptional states.

**Figure 5:**
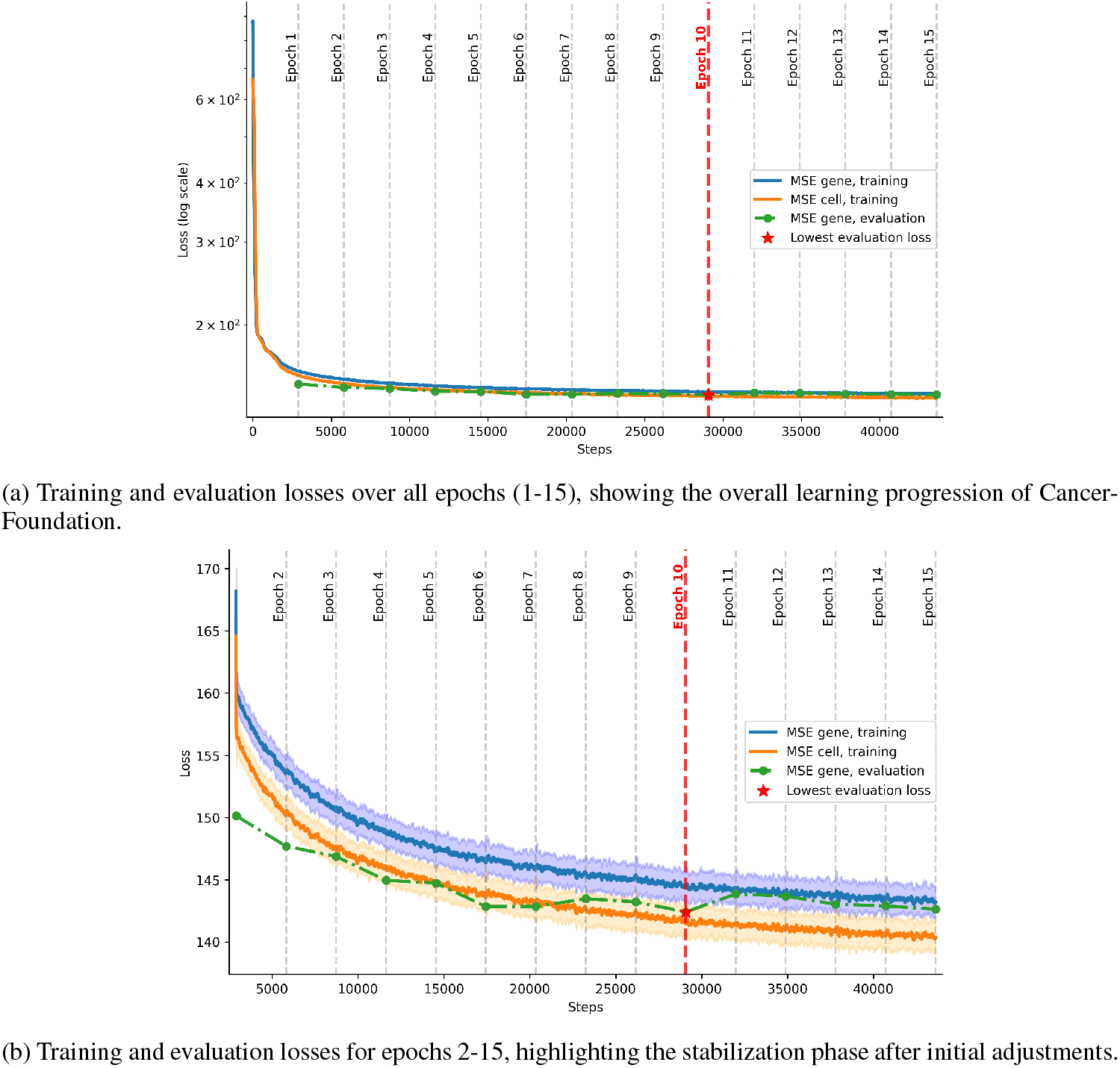
Training and evaluation losses for the pretraining of CancerFoundation. “MSE gene” refers to the loss in Equation 5, and “MSE cell” to Equation 6. Training losses are moving averages with a window size of ten, and the shaded area denotes the standard deviation. The lowest evaluation loss is marked in red.

**Figure 6:**
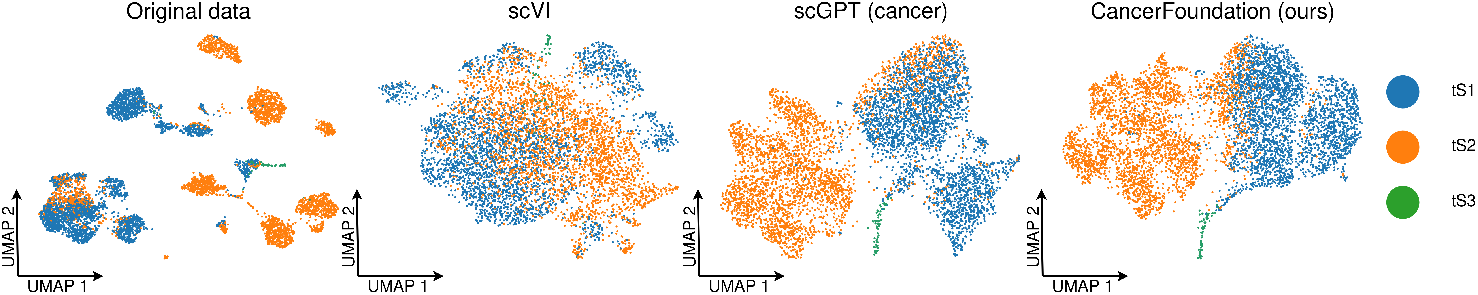
UMAP plots of cell embeddings for a lung adenocarcinoma dataset (Kim et al., 2020) colored according to the ground-truth transcriptional states: tS1 (transcriptional state 1, blue), tS2 (transcriptional state 2, yellow), tS3 (transcriptional state 3, green).

Nevertheless, the model demonstrates convergence, with the lowest evaluation loss occurring at epoch ten out of fifteen. This suggests that the model was trained for an adequate number of epochs, and extending the training further would likely not have significantly improved performance. Additionally, no signs of overfitting were observed, as the evaluation loss was consistently slightly lower than the training loss throughout.

### C Zero-shot batch integration

This appendix extends Section 4.1 by presenting additional results for batch integration performance on a different cancer type (see Appendix C.1) and analyzing the effects of our data balancing strategies (see Appendix C.2). Finally, Appendix C.3 provides a summary of key findings and concluding remarks.

#### C.1 Lung adenocarcinoma

To further assess CancerFoundation’s batch integration performance, we utilized the lung adenocarcinoma dataset from Kim et al. (2020). This dataset identified three key transcriptional states in tumor cells: tS1, tS2, and tS3. The tS1 and tS3 states represent less aggressive tumor cells that retain characteristics of normal epithelial cells, such as those involved in surfactant homeostasis and lung development. In contrast, tS2 reflects a more aggressive tumor phenotype, characterized by abnormal cell movement, proliferation, and apoptosis, and is strongly associated with late-stage cancer, metastasis, and poor patient survival.

CancerFoundation outperformed scVI and scGPT (cancer) in preserving biological signals across batches along all metrics (see Table 3). scGPT and our method, CancerFoundation, performed similarly, with CancerFoundation excelling in the isolated labels and silhouette label metric, whereas scGPT was superior for the Leiden-related scores (Traag et al., 2019). Interestingly, scGPT outperforms its cancer-specific variant, scGPT (cancer), for the Leiden-related metrics by a large margin. As for the batch correction metrics, scVI has an edge over the other methods, particularly in terms of the iLISI score.

**Table 3:** Batch integration metrics evaluating the performance of different embeddings in integrating the batches for a lung adenocarcinoma dataset (Kim et al., 2020) using the framework introduced by Luecken et al. (2022).

| Method | Biological conservation |  |  |  |  | Batch correction |  |  |  | Aggregate score |  |  |
| --- | --- | --- | --- | --- | --- | --- | --- | --- | --- | --- | --- | --- |
|  | Isolated labels | Leiden NMI | Leiden ARI | Silhouette label | cLISI | Silhouette batch | iLISI | KBET | GC | Batch correction | Biological conservation | Total |
| CancerFoundation (ours) | **0.74** | 0.50 | 0.50 | **0.63** | **1.00** | 0.90 | 0.01 | 0.21 | **0.97** | 0.52 | **0.67** | **0.61** |
| scGPT | 0.56 | **0.58** | **0.63** | 0.56 | 0.97 | **0.91** | 0.02 | **0.29** | 0.95 | 0.54 | 0.66 | **0.61** |
| scGPT (cancer) | 0.65 | 0.49 | 0.49 | 0.57 | 0.99 | 0.89 | 0.00 | 0.23 | 0.94 | 0.52 | 0.64 | 0.59 |
| scVI | 0.60 | 0.14 | 0.07 | 0.52 | 0.83 | 0.88 | **0.11** | 0.25 | 0.96 | **0.55** | 0.43 | 0.48 |

#### C.2 Ablation study

As outlined in Section 3.2.1, and 3.2.4, we perform random oversampling and domain invariant training with respect to the scRNA-seq technologies, aiming to achieve improvements for underrepresented technologies. Indeed, CancerFoundation showed superb zero-shot batch integration performance for the dataset by Neftel et al. (2019) (see Section 4.1), using the Smart-Seq2 technology which constitutes a minority in our dataset. To what extent, if any, can this be attributed to oversampling and domain-invariant training?

##### Brain-cancer setting

To address this issue, we first trained a small model exclusively on brain cancer cells, excluding multiple studies that used the Smart-Seq2 technology. Consequently, cells examined using Smart-Seq2 accounted for only 4% of all brain cancer cells. We then trained three different models: one utilizing random oversampling, another employing domain-invariant training (DIT), and a third without either of these methods.

The metrics in Table 4 show that random oversampling significantly improved the conservation of the biological signal, while the effect of DIT was less pronounced.

**Table 4:** Comparison of models trained with random oversampling (RO), domain-invariant training (DIT), and without either method (baseline) on batch integration metrics (Luecken et al., 2022), using the dataset from Neftel et al. (2019) (Smart-Seq2). The models were trained on a modified brain cancer dataset, where cells profiled using Smart-Seq2 represent only 4% of the total dataset.

| Method | Biological conservation |  |  |  |  | Batch correction |  |  |  | Aggregate score |  |  |
| --- | --- | --- | --- | --- | --- | --- | --- | --- | --- | --- | --- | --- |
|  | Isolated labels | Leiden NMI | Leiden ARI | Silhouette label | cLISI | Silhouette batch | iLISI | KBET | GC | Batch correction | Biological conservation | Total |
| RO | **0.56** | **0.45** | **0.39** | **0.56** | **0.91** | **0.80** | 0.09 | 0.38 | **0.96** | 0.56 | **0.57** | **0.52** |
| DIT | **0.56** | 0.42 | 0.28 | 0.55 | 0.90 | 0.79 | 0.10 | **0.45** | **0.96** | **0.58** | 0.54 | 0.51 |
| Baseline | 0.55 | 0.38 | 0.28 | 0.55 | 0.89 | 0.79 | **0.11** | **0.45** | **0.96** | **0.58** | 0.53 | 0.50 |

##### Pan-cancer setting

These findings prompted us to train an additional version of CancerFoundation without random oversampling or DIT, on all the data, in order to better isolate the effects of these methods. As shown in Table 5, incorporating random oversampling and DIT significantly enhances the preservation of transcriptional states. Notably, Figure 7 demonstrates that CancerFoundation, when trained without oversampling and DIT, struggles to distinguish MES-like cells from AC-like cells, similar to scGPT (cancer).

**Table 5:**
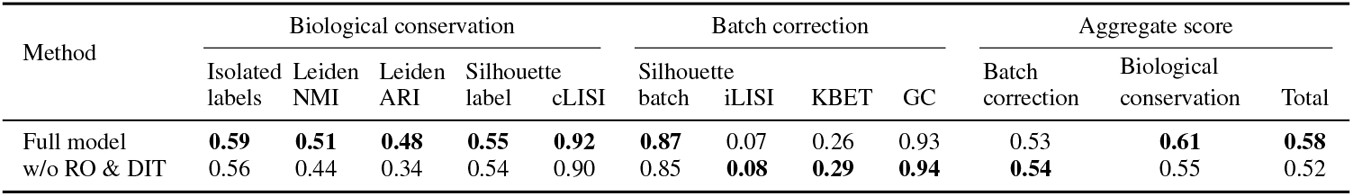
Batch integration metrics evaluating the performance of different embeddings in integrating the batches (Luecken et al., 2022) for a glioblastoma dataset (Neftel et al., 2019). Full model: CancerFoundation when employing random oversampling (RO) and domain invariant training (DIT) as in the standard setup. w/o RO & DIT: CancerFoundation without using RO and DIT.

| Method | Biological conservation |  |  |  |  | Batch correction |  |  |  | Aggregate score |  |  |
| --- | --- | --- | --- | --- | --- | --- | --- | --- | --- | --- | --- | --- |
|  | Isolated labels | Leiden NMI | Leiden ARI | Silhouette label | cLISI | Silhouette batch | iLISI | KBET | GC | Batch correction | Biological conservation | Total |
| Full model | **0.59** | **0.51** | **0.48** | **0.55** | **0.92** | **0.87** | 0.07 | 0.26 | 0.93 | 0.53 | **0.61** | **0.58** |
| w/o RO & DIT | 0.56 | 0.44 | 0.34 | 0.54 | 0.90 | 0.85 | **0.08** | **0.29** | **0.94** | **0.54** | 0.55 | 0.52 |

**Table 6:** Survival prediction performance for different embeddings and gene expression as baseline. Values show mean Harrell’s concordance index followed by standard deviation computed across 25 folds. Percentages in parentheses indicate the proportion of samples for each cancer type in the CancerFoundation training data.

| Model | KIRC (3%) | LGG (2%) | LIHC (2%) | HNSC (2%) | COAD (2%) | UCEC (1%) | SARC (1%) |
| --- | --- | --- | --- | --- | --- | --- | --- |
| CancerFoundation (ours) | 0.67±0.07 | 0.78±0.05 | 0.63±0.09 | 0.52±0.06 | 0.51±0.08 | 0.63±0.09 | 0.65±0.08 |
| scGPT (cancer) | 0.66±0.07 | 0.77±0.07 | 0.63±0.09 | 0.52±0.06 | 0.54±0.08 | 0.63±0.10 | 0.66±0.07 |
| scGPT | 0.68±0.07 | 0.77±0.04 | 0.61±0.09 | 0.54±0.07 | 0.54±0.08 | 0.64±0.08 | 0.66±0.06 |
| Baseline (GEX) | 0.70±0.05 | 0.82±0.05 | 0.65±0.08 | 0.57±0.06 | 0.56±0.07 | 0.72±0.09 | 0.63±0.07 |
|  | BLCA (0%) | CESC (0%) | ESCA (0%) | KIRP (0%) | READ (0%) | STAD (0%) |  |
| CancerFoundation (ours) | 0.59±0.04 | 0.52±0.17 | 0.48±0.07 | 0.81±0.14 | 0.51±0.15 | 0.52±0.04 |  |
| scGPT (cancer) | 0.60±0.04 | 0.60±0.16 | 0.49±0.08 | 0.82±0.14 | 0.53±0.14 | 0.53±0.05 |  |
| scGPT | 0.60±0.05 | 0.54±0.18 | 0.48±0.07 | 0.82±0.12 | 0.57±0.17 | 0.52±0.05 |  |
| Baseline (GEX) | 0.61±0.05 | 0.56±0.12 | 0.48±0.07 | 0.83±0.11 | 0.57±0.11 | 0.56±0.05 |  |

**Figure 7:**
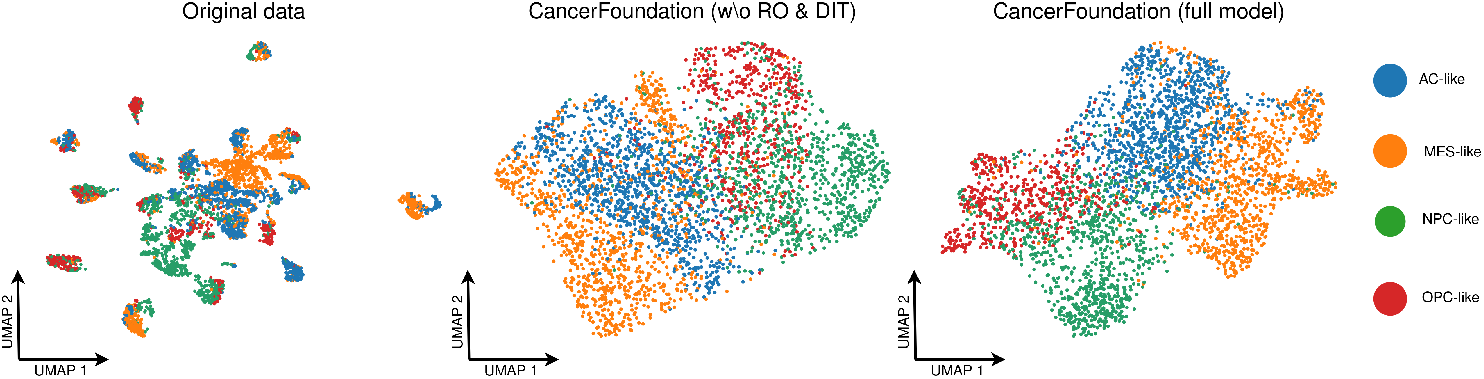
UMAP plots of cell embeddings colored according to the ground-truth transcriptional states for a glioblastoma dataset (Neftel et al., 2019): AC-like (astrocyte-like, blue), MES-like (mesenchymal-like, yellow), NPC-like (neural-progenitor-like, green), and OPC-like (oligodendrocyte-progenitor-like, red).

#### C.3 Discussion

The results for zero-shot batch integration in Section 4.1 reveal a multitude of interesting insights. For one, despite being trained on reconstructing raw data that exhibit strong patient-specific effects, CancerFoundation’s embeddings demonstrated an *emergent ability* to correct for these effects and group cells by their transcriptional state. Importantly, this zero-shot performance, achieved without any prior training on these datasets, highlights the robustness of our foundation model.

Moreover, despite being trained on five million cancer-related cells, including both malignant and non-malignant types, scGPT (cancer) (Cui et al., 2023) does not consistently outperform scGPT, which was trained on a much larger set of 30 million primarily peripheral blood mononuclear cells. In contrast, CancerFoundation, which was trained on only 1 million malignant cells, achieves superior performance. One possible explanation for the lack of advantage seen in scGPT (cancer) compared to scGPT could be that the inclusion of both malignant and non-malignant cells during training introduces greater variability and noise, diluting the model’s ability to focus on the distinct transcriptional states of malignant cells. Malignant cells often exhibit highly distinct transcriptional programs compared to non-malignant cells, and integrating both types might obscure these spe- cific malignant features, making it harder for scGPT (cancer) to effectively identify and generalize malignant cell states during batch integration tasks.

However, the improvements in zero-shot integration achieved by CancerFoundation cannot be solely attributed to training on malignant cells, as CancerFoundation differs architecturally from scGPT. As discussed in Section C.2, the enhancements observed in the dataset from Neftel et al. (2019), using Smart-Seq2 (Picelli et al., 2013), are primarily due to our approach of random oversampling of minority scRNA-seq technologies and the application of domain-invariant training.

Finally, it is important to recognize that batches are linked to individual patients, meaning that variations between batches reflect not only potential confounding factors but also patient-specific biological signals. The fact that scVI (Lopez et al., 2018) performs well in batch correction yet struggles to preserve these biological signals across both datasets suggests that it may be overcorrecting for batch effects.

These results not only demonstrate CancerFoundation’s emergent ability to integrate batches in a zero-shot setting but also provide empirical validation of its *aptness* for cancer-specific downstream tasks, where clustering by *meta-program*, rather than by batch or patient, is of fundamental importance (Marusyk et al., 2012; McGranahan & Swanton, 2015).

### D Survival analysis

#### D.1 Implementation

All Ridge-regularized Cox PH models were implemented using `glmnet` 4.1 in `R` 4.3.3 (R Core Team, 2024; Friedman et al., 2010; Simon et al., 2011).

To run `glmnet`, covariates were standardized within `glmnet`. In addition, the path was run over 100 regularization hyperparameters with a minimum ratio of 0.01 and computing the cross-validated partial likelihood via subtraction (`grouped=TRUE`) (Verweij & Van Houwelingen, 1993). Cross-validation folds for choosing the optimal regularization hyperparameter were chosen using `splitTools` 1.0.1, by stratifying the event indicator using a fixed seed across test splits for reproducibility and then passed to `glmnet` (Mayer, 2023). Predictions were made using the regularization hyperparameter that minimizes the cross-validated partial likelihood (`lambda.min`).

For calculating Harrell’s concordance index, we used the `Python` 3.8.0 implementation in `scikit-survival` 0.22.2, `concordance_index_`censored, with the default tolerance for differentiating between ties of 1e−08 (Pölsterl, 2020).

#### D.2 Dataset

Datasets were chosen following the selection criteria of Wissel et al. (2022). In brief, the authors only keep datasets containing at least 100 samples and a minimum number of events greater than or equal to max(10, 0.05*n*), where *n* indicates the sample size of a particular dataset.

The preprocessed TCGA datasets from Wissel et al. (2022) were used without modification, discarding all covariates not related to gene expression.

#### D.3 Validation

We used the cross-validation splits generated by Wissel et al. (2022) to match their datasets for validation and performance metric calculation. The authors generate 25 total splits, performing 5-fold cross-validation, and repeat it five times.

#### D.4 On the choice of concordance index

While several other concordance indices are now popular for evaluating survival models, including Uno’s concordance and Antolini’s concordance, we chose to use Harrell’s concordance for several reasons (Uno et al., 2011; Antolini et al., 2005).

First, Uno’s concordance has its own challenges, in particular how to best estimate the censoring distribution, as estimating it on the test set may overfit and estimating it on the training set may cause numerical instabilities without proper truncation (Kvamme & Borgan, 2023). In addition, Harrell’s concordance is arguably still the most commonly used concordance measure, in particular for work not focusing solely on survival analysis. Lastly, the comparability of concordance indices across models is paramount (Sonabend et al., 2022). Since our work considers only Cox PH models, we do not require comparability to more complicated models which may not be able to make predictions on the risk measure level and can thus rely on the more simple concordance index proposed by Harrell Jr et al. (1996).

## Footnotes

1 Github repository: https://github.com/BoevaLab/CancerFoundation.

1 Pytorch embedding layer.

1 Pytorch’s cosine schedule with warmup.

